# A global overview of single-cell type selectivity and pleiotropy in complex diseases and traits

**DOI:** 10.1101/2020.11.18.388488

**Authors:** Chao Xue, Lin Jiang, Qihan Long, Ying Chen, Xiangyi Li, Miaoxin Li

**Affiliations:** Zhongshan School of Medicine, Sun Yat-sen University, Guangzhou, 510080, China; Research Center of Medical Sciences, Guangdong Provincial People’s Hospital, Guangdong Academy of Medical Sciences, Guangzhou, 510080, China; Key Laboratory of Tropical Disease Control (Sun Yat-sen University), Ministry of Education, Guangzhou, 510080, China; Center for Precision Medicine, Sun Yat-sen University, Guangzhou, 510080, China; State Key Laboratory of Brain and Cognitive Sciences, the University of Hong Kong, Pokfulam, Hong Kong

**Keywords:** single-cell, cell-type selectivity, cell-type pleiotropy, selective expression, associated gene, genome-wide association study

## Abstract

After centuries of genetic studies, one of the most fundamental questions, i.e. in what cell types do DNA mutations regulate a phenotype, remains unanswered for most complex phenotypes. The current availability of hundreds of genome-wide association studies (GWASs) and single-cell RNA sequencing (scRNA-seq) of millions of cells provides a unique opportunity to address the question. In the present study, we firstly constructed an association landscape between over 20,000 single cell clusters and 997 complex phenotypes by a cross annotation framework with scRNA-seq expression profiles and GWAS summary statistics. We then performed an extensive overview of cell-type specificity and pleiotropy in human phenotypes and found most phenotypes (>90%) were moderately selectively associated with a limited number of cell types while a small fraction cell types (<10%) had strong pleiotropy in multiple phenotypes (~100). Moreover, we identified three cell type-phenotype mutual pleiotropy blocks in the landscape. The application of the single cell type-phenotype cross annotation framework (named SPA) also explained the T cell biased lymphopenia and suggested important supporting genes in severe COVID-19 from human genetics angle. All the cell type-phenotype association results can be queried and visualized at http://pmglab.top/spa.

---

Tissue-specific manifestation is prevalent among complex diseases and traits^1^. Understanding of the cellular specificity is indispensable for unravelling their pathological mechanisms and developing precision therapy^2,3^. Identifying critical cells and genes that determine the tissue-selective pathogenesis could provide a primary step towards elucidating disease etiologies, which can help to understand resilience of unaffected tissues and further open new therapeutic directions^1^. However, probably due to lack of effective methods and data, deciphering cellular spectrum and pathological mechanisms of many complex phenotypes (diseases and traits) is lagging far behind their genome-wide association studies even for well-studied diseases, including schizophrenia^4^, arthritis^5^ and type 2 diabetes^6^. Recently, based on genes’ selective expression in bulk cells, summary statistics of genome-wide association studies (GWASs) of the complex phenotypes were utilized to explore the tissue specificity of the corresponding phenotypes^7,8^. These studies suggested that genes’ selective expression leveraged with GWAS signals may be useful for inferring cellular spectrum of many complex phenotypes.

The single-cell RNA sequencing (scRNA-seq) technology provides excellent advantages to precisely profile genes’ selective expression in cell types and deeply understand cell lineage. Nowadays, scRNA-seq technology has been widely used to detect heterogeneity among tumor cells^9,10^, to reveal developmental processes and cell fate decisions^11^, and to profile lineages and cell types in the vertebrate brain^12^. Watanabe et al. testified the cell-type selective expression of genes associated with phenotypes by using scRNA-seq data as well^13^. Recently, a human cell atlas project has provided a comprehensive human cell landscape which released gene expression profiles and cell hierarchy for over a half million cells^14^. Meanwhile, several comprehensive public resources are also available to query gene expression in single cells. For instance, the PanglaoDB to date has collected gene expression profiles for over 1 million human cells and around 4.5 million mouse cells^15^. These resources provide a unique opportunity for addressing the question of cell-type selectivity and even pleiotropy in complex diseases and traits.

To precisely and compressively understand tissue specificity and pleiotropy of complex phenotypes, we constructed a landscape of association between thousands of human single-cell clusters and around 1000 complex diseases and traits by a <u>s</u>ingle-cell type and <u>p</u>henotype cross <u>a</u>nnotation framework (named SPA). The landscape led to a systematic analysis on patterns of single-cell type selectivity and pleiotropy in complex diseases and traits. We subsequently replicated the association patterns in an independent human scRNA-seq dataset and a mouse scRNA-seq dataset respectively. Based on the association landscape, we further developed two application functions on SPA, i.e., identifying phenotype-relevant single-cell types and annotating relevant phenotype for customized single-cell clusters. Finally, we demonstrated the two functions in a scRNA-seq dataset of human lung and a GWAS summary statistics dataset of severe COVID-19 as proof-of-principle examples respectively.

## Results

### Overview of the single-cell type and phenotype cross annotation framework constructing the association landscapes and its application functions

We constructed a compressive landscape of associations between thousands of single cell clusters and about 1,000 phenotypes by a single-cell type and phenotype cross annotation framework (named SPA) according to a workflow shown in **Fig. 1b**. Firstly, we curated a huge amount of human scRNA-seq expression dataset from PanglaoDB^15^ and generated selective expression profiles of genes among thousands of single cell clusters by a robust-regression z-score approach (REZ)^8^. Secondly, we initially collected 1,871 GWAS datasets of different phenotypes with large sample sizes (n≥ 10,000). The two types of data were then analyzed by an iterative prioritization procedure. In the procedure, phenotype-associated genes were prioritized by a conditional gene-based association according to the genes’ selective expression in disease related cell-types while the phenotype related cell-types were prioritized by an enrichment analysis of Wilcoxon rank-sum test for phenotype-associated genes’ selective expression. The phenotype-associated gene list and phenotype related cell-type list were updated by turns until the two list were unchanged. The underlying assumption was that the disease susceptibility genes tend to selectively expressed in disease relevant cell-types. The iterative prioritization procedure was extended from our recent method^8^. The major extension was that a more robust conditional gene-based association test was proposed for the procedure (See details in the method section). The iterative prioritization procedure was carried out at all the phenotypes one by one and produced significant cell clusters of each phenotype and the supporting genes which were associated with the phenotype and selectively expressed in the significant cell clusters. After removing phenotypes with small number of significant genes (<40), a landscape of associations between numerous cell clusters and 997 phenotypes was obtained.

**Fig. 1:**
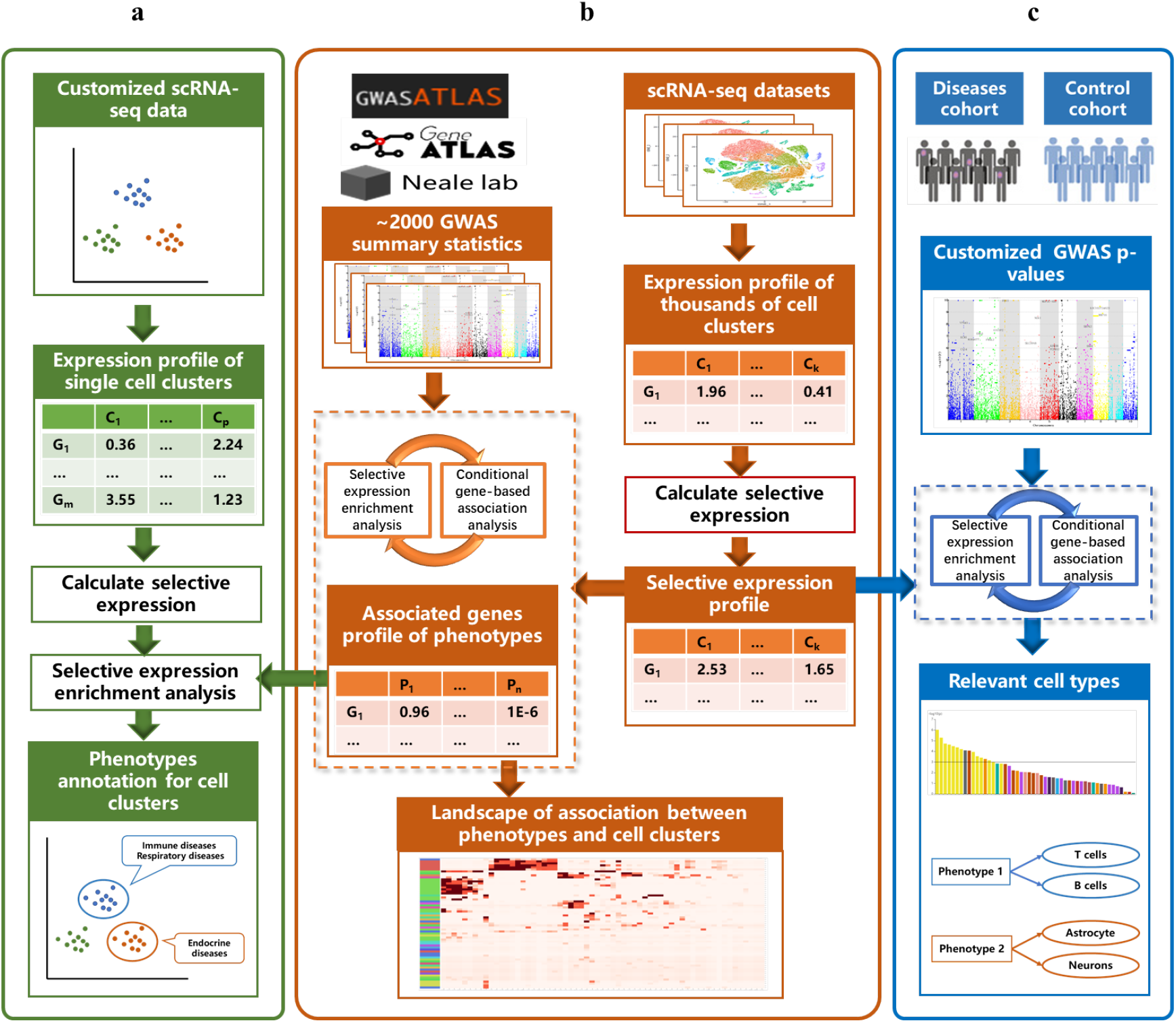
Overview of the single-cell type and phenotype cross annotation framework. **a**, Workflow of annotating customized single cell clusters with phenotypes. **b**, Workflow of constructing association landscape between around one thousand phenotypes and thousands of single cell clusters. **c**, Workflow of prioritizing the relevant cell clusters by GWAS p-values of customized phenotype.

Moreover, based on the landscape, SPA also has two types of application functions (**Fig. 1a, c**). The first application function is to annotate customized single cell clusters by complex phenotype, which can prioritize relevant phenotypes (among the 997 ones) of a cell cluster. The application function firstly calculates the selective gene expression profiles of the users’ customized single cell clusters. Then the association between customized cell clusters and around 1,000 phenotypes in the landscape are examined by the Wilcoxon rank-sum test. Contrary to the first application function, the second application function is to annotate a complex phenotype with relevant single cells, in which users can prioritize the genetically relevant cells of users’ customized phenotypes. The application function is the same as above iterative prioritization procedure in which the selective expression profiles of thousands of cell clusters in the landscape are used. We took a scRNA-seq dataset of human lung and a GWAS summary statistics dataset of severe COVID-19 as proof-of-principle examples to show the effectiveness of the two application functions respectively.

We developed a friendly web server, available at http://pmglab.top/spa, for users to query all associated cell clusters of a complex phenotype and the vice versa in the landscape (**Supplementary Fig. S4**). The two application functions of the framework can also be carried out on the web server dynamically. All involved resource data in the framework can be directly downloaded from the web server, including associated genes profiles of phenotypes, gene expression profiles of cell clusters and the entire association landscape.

### Selective expression profiles in single cell clusters

We collected gene expression profile of 1,126,580 human single cells in 3,816 clusters from PanglaoDB^15^ and calculated average expression of the cell clusters. To evaluate the similarity of the cell clusters derived from different studies, we projected the genes expression of them into a two-dimensional map using t-SNE^16^ (**Fig. 2a-b**). Most single cell clusters were grouped together according to their origins despite they were from different datasets. For example, the clusters inferred as reproductive cells were grouped together while they were from different datasets, SRA645804, SRA667709, and SRA826293.

**Fig. 2:**
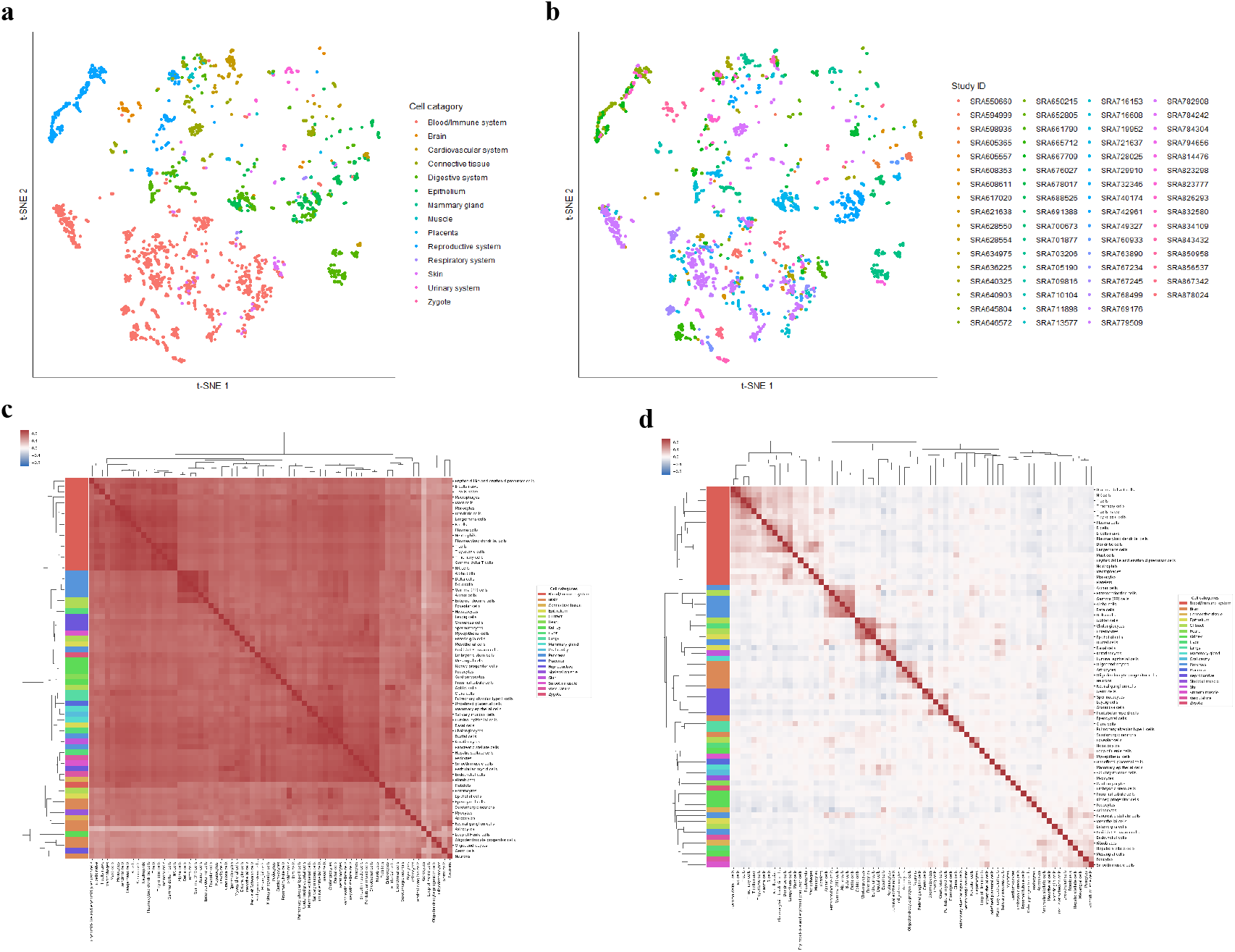
Average expression profile and selective expression profile of single cell clusters. **a-b,** Cell clusters visualization by t-SNE dimensionality reduction according to averaged expression profiles. Each data point represents a cell cluster. There are 2,305 clusters after excluding the clusters inferred as unknown cell types. They are colored by (**a**) categories of inferred cell type according to anatomy and (**b**) NCBI SRA dataset ID. **c-d**, Pearson correlation coefficient of the cell types according to (**c**) original expression profiles and (**d**) selective expression profiles. Cell types are clustered based on the pair-wise correlation matrix using the hierarchical clustering. The color bar at the right of the heatmap represents the categories of cell types according to anatomy.

Next, we calculated the selective expression profiles of the clusters by a robust-regression z-score method (REZ) that we previously proposed^8^. To check whether the usage of selective expression can improve discrimination between cell clusters, we investigated the Pearson correlation coefficient of inferred cell types according to original expression and selective expression respectively (**Fig. 2c-d**). As expected, the correlation coefficient of same cell types was higher than that of different ones according to both types of expression profiles, such as the immune/blood cells and pancreas cells. This was also true for the cell types of similar origins, such as enterocyte and epithelial cells with Pearson correlation coefficients 0.98 and 0.86 in original expression and selective expression profiles respectively. However, when the cell types were different, based on original expression profiles, the correlation coefficients were still relatively high, such as enterocyte and macrophages with Pearson correlation coefficient 0.71. Probably, the high correlation coefficient was because of expression of house-keeping genes. In contrast, based on the selective expression profile, the correlation coefficient between clusters from different cell-types became very low (~0). Therefore, our result suggested the selective expression was more effective than the original expression to discriminate single-cell clusters, which was consistent with our previous findings on the bulk RNA expression^8^.

### Associated genes profile of massive traits and diseases

After quality control in 2250 GWAS datasets, we had full GWAS summary statistics of 1871 traits and diseases from three public databases (See details in the methods section). We then performed an iterative conditional gene-based analysis by our proposed effective chi-square test (See it in methods section) with the above selective expression profiles in single cell clusters. Finally, 997 phenotypes with more than 40 conditionally significant associated genes were obtained for next curation and annotation (**Supplementary Fig. S1**). We manually assigned the 997 phenotypes to 19 categories according to their clinical characteristics. It should be noted that 366 out of 997 phenotypes were assigned in to “other” category due to ambiguous clinical characteristics (**Supplementary Table S1**).

We further checked the similarity between different phenotypes according to their overlapped significant genes in the 18 clinical categories (excluding the category “other”). The similarity was measured by average Jaccard coefficient based on the associated genes (**Fig. 3a**). Sixteen phenotype categories had higher similarity with themselves than that with other categories, which suggested a general consistency between clinical definition and genetic similarity of the phenotypes. However, we also noticed that some distinct clinical categories had relatively high genetic similarity, such as metabolic phenotypes vs. body measure phenotypes and endocrine phenotypes vs. respiratory phenotypes. We further calculated the Jaccard similarity of phenotypes belonging to the endocrine and respiratory categories according to their associated genes (**Fig. 3b**). Some phenotypes belonging to different categories had high similarity, such as thyrotoxicosis (belonging to endocrine phenotypes) and asthma (belonging to respiratory phenotypes). This is probably because both diseases were related to autoimmunity^17,18^. We also noticed some phenotypes within endocrine diseases had low similarity, such as thyrotoxicosis and adrenal disorder. Therefore, we further clustered the phenotypes according to the phenotype-associated genes within each clinical category by hierarchical clustering (see details in methods section). The 997 phenotypes were classified into 107 sub-categories finally (**Fig. 6c, Supplementary Table S1**). The sub-categories are named by combining the clinical category name and cluster ID. For example, the phenotype cluster “1” in “respiratory” category is named as ‘respiratory_’.

**Fig. 3:**
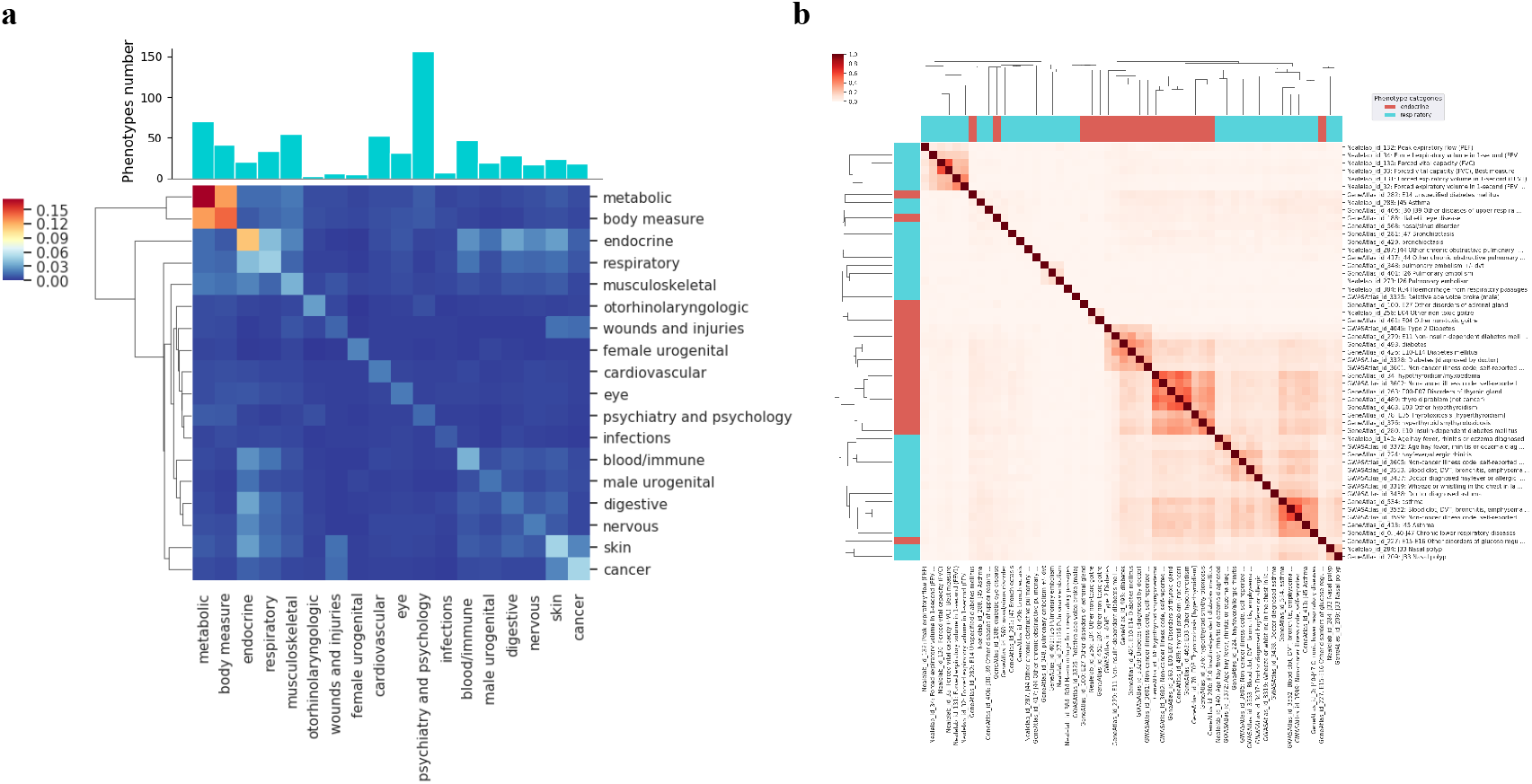
Characterization of associated genes profile of phenotypes. **a**, The top bar plot shows the phenotypes number of different categories. The heatmap shows the average Jaccard similarity coefficient among the phenotype categories according to the associated genes (See details in methods section). The categories are clustered based on the pair-wise correlation matrix using the hierarchical clustering. The category “other” was excluded from figure due to the mixture of its phenotypes. **b**, Jaccard similarity of phenotypes belonging to endocrine, respiratory phenotypes. Phenotypes are clustered based on the pair-wise correlation matrix using the hierarchical clustering. The color bar at the top-right of the heatmap represents the clinical categories of phenotypes.

**Fig. 4:**
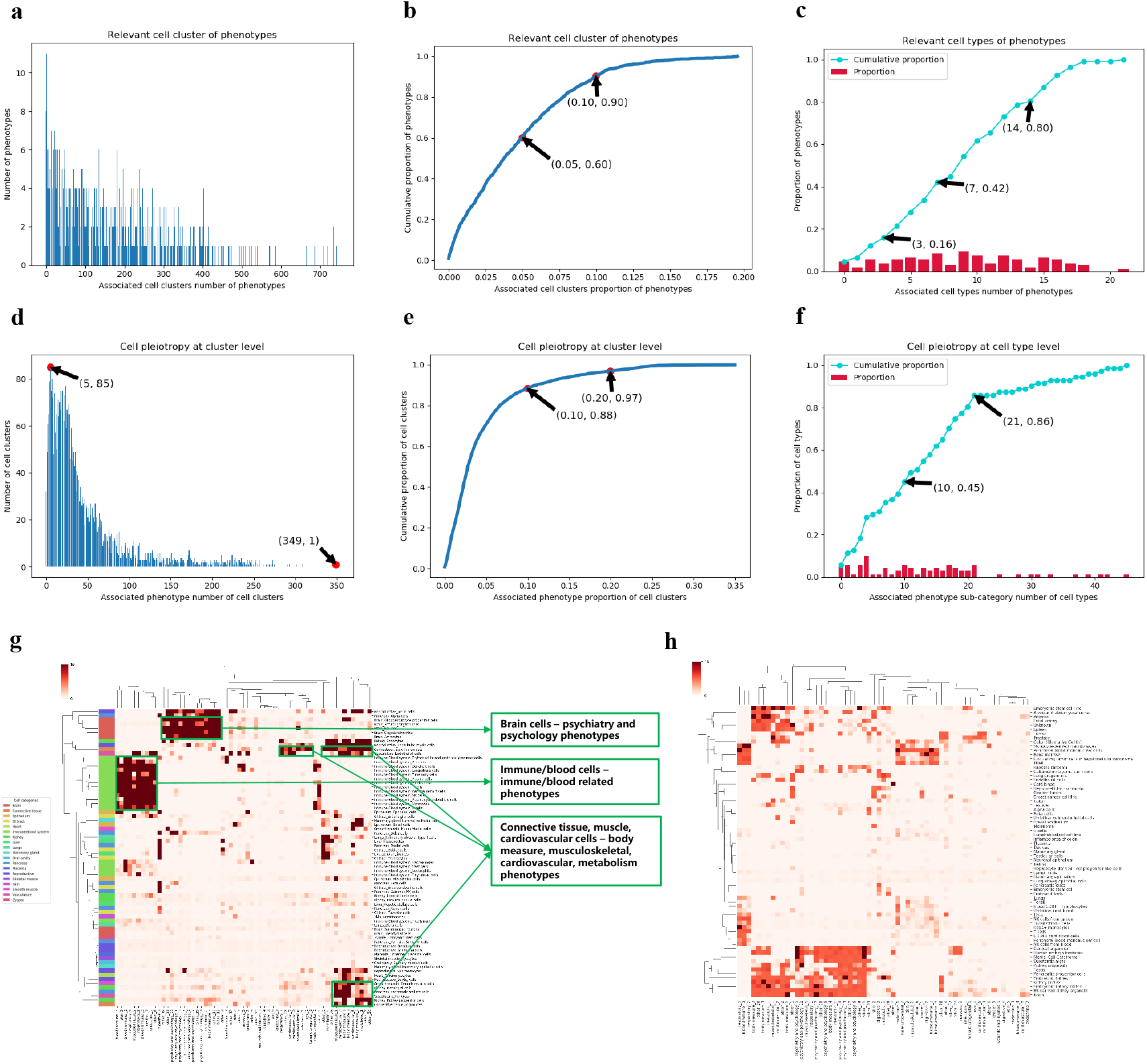
Association landscape between single cell clusters and phenotypes. **a**, The histogram shows the frequency of phenotypes with different number of associated cell clusters. **b**, The curve shows the accumulative proportion of phenotypes with the different proportion of associated cell clusters. **c**, The height of the red bar shows the proportion of phenotype sub-categories with different number of associated cell types and the blue curve shows the accumulative proportion. **d**, The histogram shows the frequency of cell clusters with different number of associated phenotypes. **e**, The curve shows the accumulative proportion of cell clusters with the different proportion of associated phenotypes. **f**, The height of the red bar shows the proportion of cell types with different number of associated phenotype sub-categories and the blue curve shows the accumulative proportion. **g**, Enrichment of cell types in the phenotype sub-categories. The color bar at the left of the heatmap represents the categories of cell types according to anatomy. **h**, Enrichment of source tissues/cells of clusters inferred as unknown cell types in the phenotype sub-categories. **g-h**, The heatmaps are colored by the −log10(p) of the hypergeometric test. Both of two axes are clustered based on the pair-wise correlation matrix using the hierarchical clustering. For clarity and convenience, we only showed the phenotype sub-categories with no fewer than five phenotypes.

**Fig. 5:**
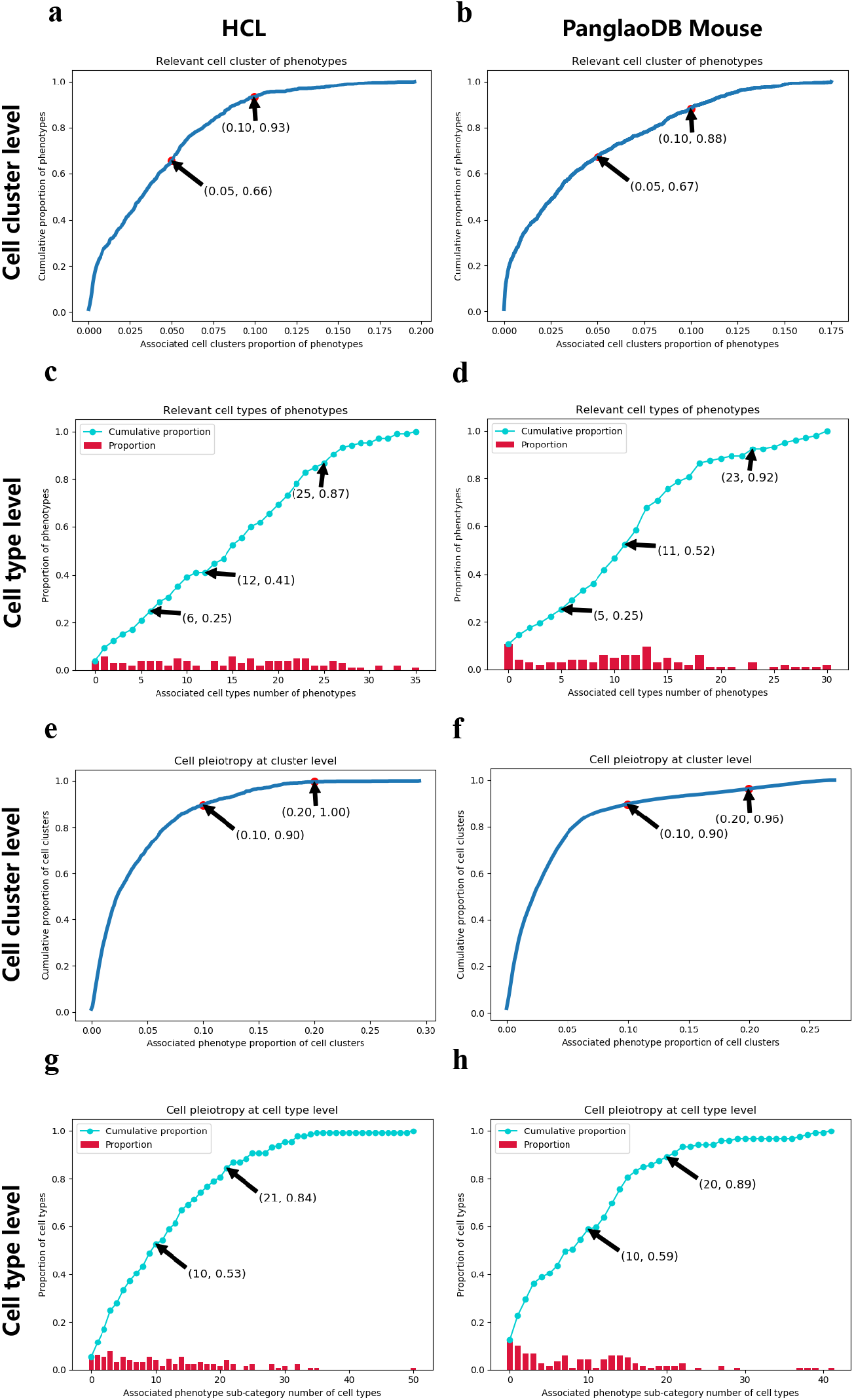
Validation with an independent scRNA dataset. **a-b**, The curve shows the accumulative proportion of phenotypes with different proportion of associated cell clusters in (**a**) HCL and (**b**) PanglaoDB Mouse dataset respectively. **c-d**, The height of red bar shows the proportion of sub-categories with different number of associated cell types and the blue curve shows the accumulative proportion in (**c**) HCL and (**d**) PanglaoDB Mouse dataset respectively. **e-f**, The curve shows the accumulative proportion of cell clusters with different proportion of associated phenotypes in (**e**) HCL and (**f**) PanglaoDB Mouse dataset respectively. **g-h**, The height of red bar shows the proportion of cell types with different number of associated phenotype sub-categories and the blue curve shows the accumulative proportion in (**g**) HCL and (**h**) PanglaoDB Mouse dataset respectively.

**Fig. 6:**
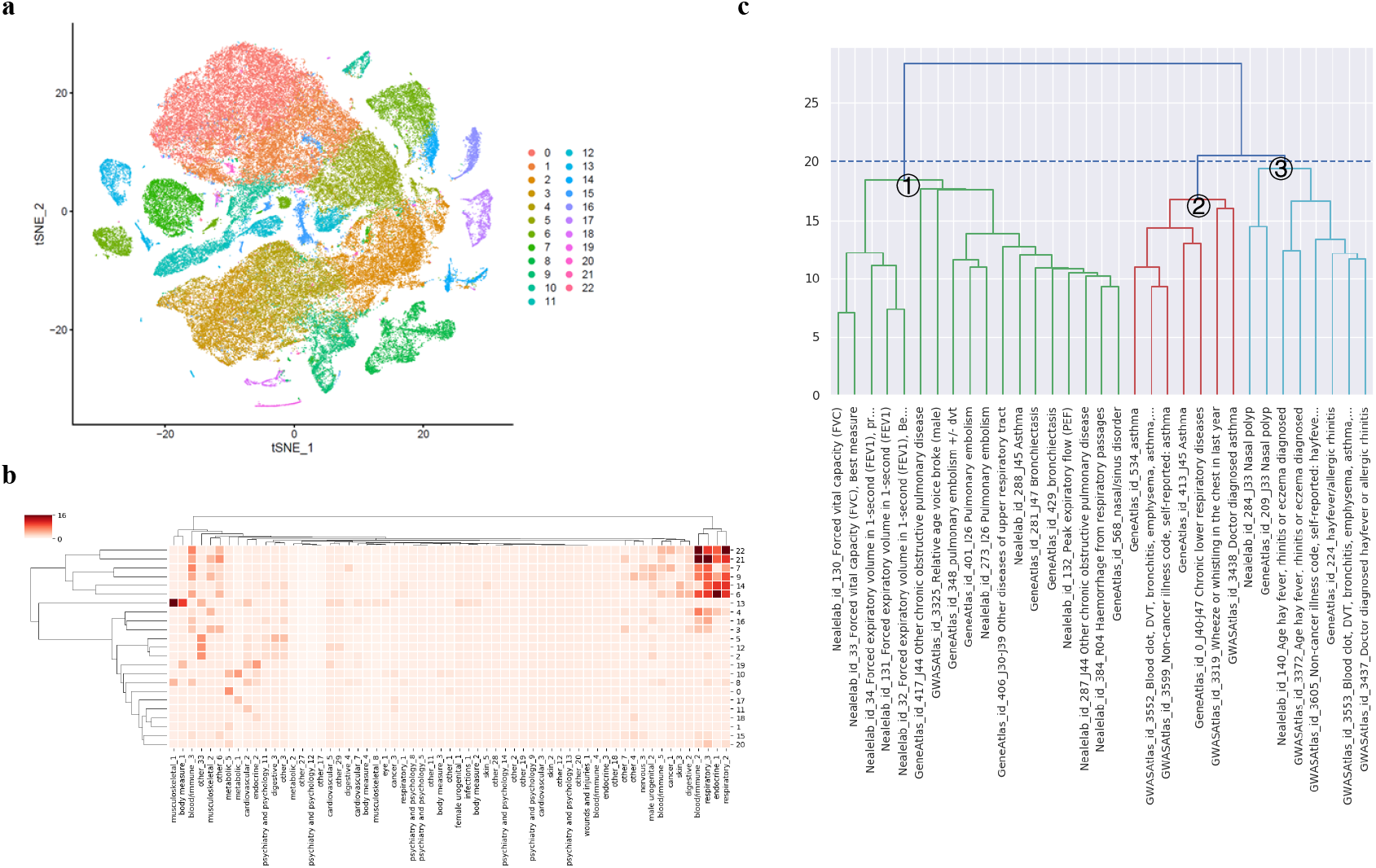
Phenotypes annotation of the single-cell clusters from lung. **a,** t-SNE visualization of 91,914 single cells from 8 lung samples of human. The cells were clustered into 23 cell clusters labeled as 0-22. **b**, Phenotype annotation of the 23 cell clusters. The cell clusters were enriched into the phenotype sub-categories. The heatmap is colored by the −log10(p) of the hypergeometric test. Both of two axes are clustered based on the pair-wise correlation matrix using the hierarchical clustering. **c,** Hierarchical clustering of respiratory phenotypes according to the associated genes. The number 1-3 shows the phenotype cluster ID. The sub-categories are named by combining the clinical category name and cluster ID. For example, the cluster 1 is named as ‘respiratory_?. The blue horizontal line shows the distance threshold for hierarchical clustering.

### Overview of the association landscape between single cell clusters and phenotypes

We then generated a landscape of association between the above 3816 human cell clusters and 997 phenotypes. We found 175,804 (4.6%) cell cluster - phenotype pairs were significantly associated (FDR<0.01) out of 3,804,552 possible pairs. In addition, the cell-type selectivity of phenotypes and cell-type pleiotropy in phenotypes were investigated on the landscape. As expected, most phenotypes are cell-type specific or selective. The result showed about 90% phenotypes were associated with less than 10% cell clusters and even 60% phenotypes were associated with less than 5% cell clusters (**Fig. 4a-b**). On the other hand, we observed a small fraction of single cell clusters were pleiotropic in phenotypes. For example, about 12% cell clusters were associated with more than 10% phenotypes and only 3% cell clusters were associated with more than 20% phenotypes (**Fig. 4d-e**).

Due to the unbalanced distribution of cell clusters and phenotypes, we also investigated the association at cell types and phenotypes sub-categories level by enrichment of the cell types in the above 107 phenotype sub-categories (see details in methods section). In all of the 3,816 cell clusters, 2,305 clusters had known inferred cell types while 1,511 ones had unknown inferred cell types. The 2,305 clusters are inferred as 71 known cell types ranging the whole body. Similar to the above association results at cell cluster level, phenotypes sub-categories were selectively associated with a limited number of cell types. About 80% phenotypes sub-categories were associated with less than 20% cell types (**Fig. 4c**). However, we also observed that a small fraction of phenotype sub-categories were associated with many cell types. For example, the top 3 phenotype sub-categories with most associated cell types were “musculoskeletal_1” (associated with 21 (~30%) cell types, “respiratory_3” (associated with 18 (~25%) cell types) and “digestive_2” (associated with 18 (~25%) cell types) respectively (**Supplementary Table S2**). For cell type pleiotropy, we found only 14% cell types were significantly associated with more than 20% phenotype sub-categories (**Fig. 4f**). The top 3 of the most pleiotropic cell types were germ cells, neurons and fibroblasts, which were associated with 45 (42%), 42 (39%), 41 (38%) phenotype sub-categories respectively (**Supplementary Table S3**).

We further found significant association signals fell in three blocks (**Fig. 4g**), i.e. (1) brain cells with several psychiatry and psychology phenotypes, (2) immune/blood cells with immune/blood phenotypes, immune related respiratory phenotypes and endocrine phenotypes, (3) connective tissue cells, muscle cells, cardiovascular cells with body measure, musculoskeletal, cardiovascular, metabolism phenotypes. For example, the NK cells were enriched in “blood/immune_2”, “blood/immune_3”, “endocrine_1” (mainly contains thyrotoxicosis, see details in **Supplementary Table S1**), “respiratory_2” (mainly contains asthma), “respiratory_3” (mainly contains rhinitis, fever) and so on with p<1E-16. According to known knowledge, these cells play an important role in the development of these traits/diseases^17,19–27^, which suggested the associations are reliable and biologically significant. As a block means that a cell-type is associated multiple phenotypes and a phenotype is associated multiple cell-types, the block is called cell-type and phenotype mutual pleiotropy (CPMP) blocks throughout the paper.

To evaluate the associated phenotypes of cell clusters with unknown cell types, we performed the enrichment analysis of source tissues/cells of the clusters in the phenotype sub-categories (**Fig. 4h**). Similarly, we also saw that two major CPMP blocks. These unknown clusters sampled from some immune/blood tissues/cells (peripheral blood mononuclear cells, bone marrow, NK cells, T cells) were significantly enriched into blood/immune and respiratory phenotypes sub-categories. These unknown cell clusters sampled from brain tissues (human embryo forebrain, cortical organoids, substantia nigra) were enriched into the several psychiatry and psychology phenotypes sub-categories. The consistence between source tissues/cells and phenotypes also suggested the reliability of the association results.

### Validation in an independent human scRNA-seq dataset

To validate the robustness of the framework and the above observed cell-type selectivity of phenotypes and pleiotropy, we generated another association landscape with 1,606 single-cell clusters from another independent scRNA-seq dataset of human, Human Cell Landscape (HCL)^28^. Similarly, most (93%) phenotypes were selectively associated with less than 10% cell clusters (**Fig. 5a**). A smaller fraction of cell clusters were pleiotropic. Specifically, ten percent cell clusters were associated with more than 10% phenotypes and even only less than 1% cell clusters were associated with more than 20% phenotypes (**Fig. 5e**).

With regard to cell types – phenotype sub-categories association, we found 87% phenotype sub-categories were associated with less than 20% cell types (**Fig. 5c**). About 16% cell types were pleiotropic with more than 20% associated phenotype sub-categories (**Fig. 5g**). The mainly significant enrichment also fell into same CPMP blocks, i.e., (1) immune/blood cells – immune/blood phenotypes, (2) brain cells – psychiatry and psychology phenotypes, (3) connective tissue cells, muscle cells, cardiovascular cells - body measure, musculoskeletal, cardiovascular, metabolism phenotypes (**Supplementary Fig. S2a**), which were consistent with the patterns in above PanglaoDB dataset. Besides, quite a number of significant enrichments can be explained by known pathology. For example, two gastrointestinal cell types, i.e. goblet cell and enterocyte, were significantly associated with “digestive_3” (p=3.47E-9, 1.12E-9 respectively). The successfully replicated association patterns in the landscape by the independent dataset suggested the reliability of the annotation framework and our findings in cell-type selectivity of phenotypes and pleiotropy.

### Validation with a mouse scRNA-seq dataset

We then asked whether the robustness in above association patterns also conserved across species, which may help to prioritize proxy cell types in the widely-used animal models for human diseases/traits research. The validation was carried out in 16,244 single-cell clusters from mouse samples of PanglaoDB. Similar to the human datasets, most cell clusters were selectively associated with a limited number of phenotypes while a paucity of cell clusters showed strong pleiotropy. At cell cluster level, about 88% phenotypes were associated with less than 10% cell clusters (**Fig. 5b**) and only 10% cell clusters were pleiotropic with more than 10% associated phenotypes (**Fig. 5f**). At cell type level, about 92% phenotype sub-categories were associated with less than 23 (about 20%) cell types (**Fig. 5d**). For cell type pleiotropy, about 11% cell types were associated with more than 20 (about 20%) phenotypes sub-categories (**Fig. 5h**).

Similarly, the three main CPMP blocks in human datasets also appeared in the mouse dataset (**Supplementary Fig. S2b**). For example, the brain pyramidal cells was significantly enriched in “psychiatry and psychology_3”, “psychiatry and psychology_5”, “psychiatry and psychology_6”, “psychiatry and psychology_7”, etc. (p<1E-16) (**Supplementary Table S8**). The consistent association results between mouse and human scRNA datasets suggested the high conservation of gene selective expression of cells between the two different species. Although the significance level of the CPMP blocks based on the mouse cells was generally smaller than that based on the human cells, the general consistency of annotation results between the two species suggested the framework was useful for finding out relevant cells for some human diseases/traits in the mouse models.

### Application to annotate single-cell clusters from lung with associated phenotypes

We took an independent scRNA dataset of lung as proof-of-principle examples to show the usage of the customized phenotype annotation application in SPA. We collected expression profiles of 91,914 single cells from 8 human lungs^29^. The cells were clustered into 23 cell clusters by the tool Seurat^30,31^ (**Fig. 6a**). Then we annotated the 23 cell clusters with the above phenotypes by SPA and enriched the cell clusters in phenotype sub-categories (**Fig. 6b**). The main significant associations fell in six single-cell clusters, i.e. “22”, “21”, “14”, “9”, “7” and “6”, which were enriched into “respiratory_2” (mainly contains asthma), “respiratory_3” (mainly contains rhinitis, fever), “blood/immune_2” (mainly contains lymphocyte count/percent) and “endocrine_1” (mainly contains thyrotoxicosis, see details in **Supplementary Table S1**). For example, the cluster “22” was significantly enriched in “respiratory_2” and “respiratory_3” with p values 1E-16 and 5.11E-11 respectively. The significant annotations also contained many immune related phenotypes, which suggested main diseases of lung may be related with immune dysfunction and the above six cell clusters may play a key role in the pathology.

### Application to annotate severe COVID-19 with the genetically relevant cells

COVID-19 is raging around the world recently and it remains unknown which cell types are most relevant, which are important for the understanding of pathology and effective treatment. By using SPA, we prioritized the single-cell clusters related to severe COVID-19 with summary statistics of a recent GWAS^32^. To include more genes for the enrichment analysis in the small-scaled GWAS (1,610 cases vs. 2,205 controls), looser thresholds (i.e. FDR 0.25 and 10kb gene boundary extension) were used in the conditional gene-based association analysis, in which 26 significantly associated genes were obtained (**Supplementary Table S4**). The subsequent enrichment analyses were performed in two independent reference human scRNA datasets of the landscape (curated from PanglaoDB and HCL respectively). The results showed that the top five significantly associated cell clusters of severe COVID-19 in both reference datasets fell into the immune/blood system (**Table 1**). Neutrophils was prioritized as the most significant cells (p=3.5E-4) based on the dataset from PanglaoDB, while hematopoietic stem and progenitor cells was prioritized as the most significant cells based on the dataset from HCL, p=3.6E-4. Notably, there were three and two T cell clusters among the top five significant cell clusters in PanglaoDB and HCL respectively, which suggested the T cells may also play a key role in the development of severe COVID-19. Present studies showed that lymphopenia is a prominent feature of COVID-19 and seems to be more selective for T cell lineages^33^. Moreover, a recent preprint paper suggested that virus-specific TH1 and CD8+ T cells were minimally induced in patients with severe COVID-19. In contrast, asymptomatic or mildly symptomatic patients mounted potent virus-specific TH1 and CD8+ T cell responses^34^. Our analysis further conformed this finding. More importantly, our prioritized candidate susceptibility genes selectively expressed in T cells may be responsible for the weak T cell response in severe patients. For instance, we found a long intergenic non-protein coding gene, LINC00861, was not only significantly associated with COVID-19 (p=1.28E-4), but also selectively expressed in the T cell clusters (**Supplementary Table S4**), which maybe a key gene in the development of severe COVID-19. However, the function of LINC00861 is unclear at present and needs further verification by wet experiments.

**Table 1.** Top five genetically relevant cell clusters of severe COVID-19.

| Cluster ID <sup>a</sup> | P value <sup>b</sup> | Cell type <sup>c</sup> |
| --- | --- | --- |
| ***PanglaoDB*** |  |  |
| SRA779509_SRS3805266.14 | 0.00035 | Neutrophils |
| SRA550660_SRS2089638.3 | 0.0015 | Gamma delta T cells |
| SRA779509_SRS3805249.9 | 0.0023 | Erythroid-like and erythroid precursor cells |
| SRA779509_SRS3805258.5 | 0.0025 | Gamma delta T cells |
| SRA814476_SRS4073850.3 | 0.0027 | T cells |
| ***HCL*** |  |  |
| CordBlood2_5 | 0.00036 | Hematopoietic Stem and Progenitor Cells |
| CordBloodCD34P1_1 | 0.0017 | Hematopoietic Stem and Progenitor Cells |
| AdultLiver4_5 | 0.0022 | Activated T cell |
| AdultThyroid2_1 | 0.0029 | T cell |
| CordBloodCD34P2_1 | 0.0042 | Hematopoietic Stem and Progenitor Cells |

<sup>a</sup> ID of single cell cluster.
<sup>b</sup> p value of enrichment test.
<sup>c</sup> inferred cell type of cell cluster.

### Type 1 error and power of the new conditional gene-based test

In addition, we also investigated the type 1 error and power of the new conditional gene-based association test by intensive computer simulations, a key component of our framework, SPA, for generating credible phenotype-association gene profiles. The basic function of the conditional gene-based test is to remove redundant association among genes because of LD. Compared with our previous conditional gene-based test, named effective chi-square statistic (ECS)^35^, the new one does not need empirical inflation factor adjustment because it can more accurately model the dependency in marginal chi-square statistic under alterative hypothesis (See detail in method section). As shown in **Fig. 7**, the improved conditional gene-based test effectively removed redundant association from nearby genes. In six different scenarios (See detail in method section), the conditional p-values of the genes without true casual loci approximately followed uniform distribution U[0,1] regardless of the variance explained by its nearby genes because of LD and gene sizes. These results suggested the improved ECS without inflation factor adjustment could produce more valid p-values for statistical inference. In contrast, the unconditional association test produced inflated p-value based on the same gene due to its LD with nearby causal genes. With regard to statistical power, the improved ECS had slightly higher power than likelihood ratio test based on linear regression model. As shown in **Supplementary Fig. S3**, the conditional association test produced smaller p-values than the likelihood ratio test at all the scenarios regardless of the gene sizes and heritability. Noted another advantage of the improved ECS over the likelihood ratio test was that the former did not require raw genotypes of samples. The reason why the improved ECS method without using raw genotypes is still more powerful than the likelihood ratio test may be that the degree of freedom in the latter is inflated by the LD among variants.

**Figure 7:**
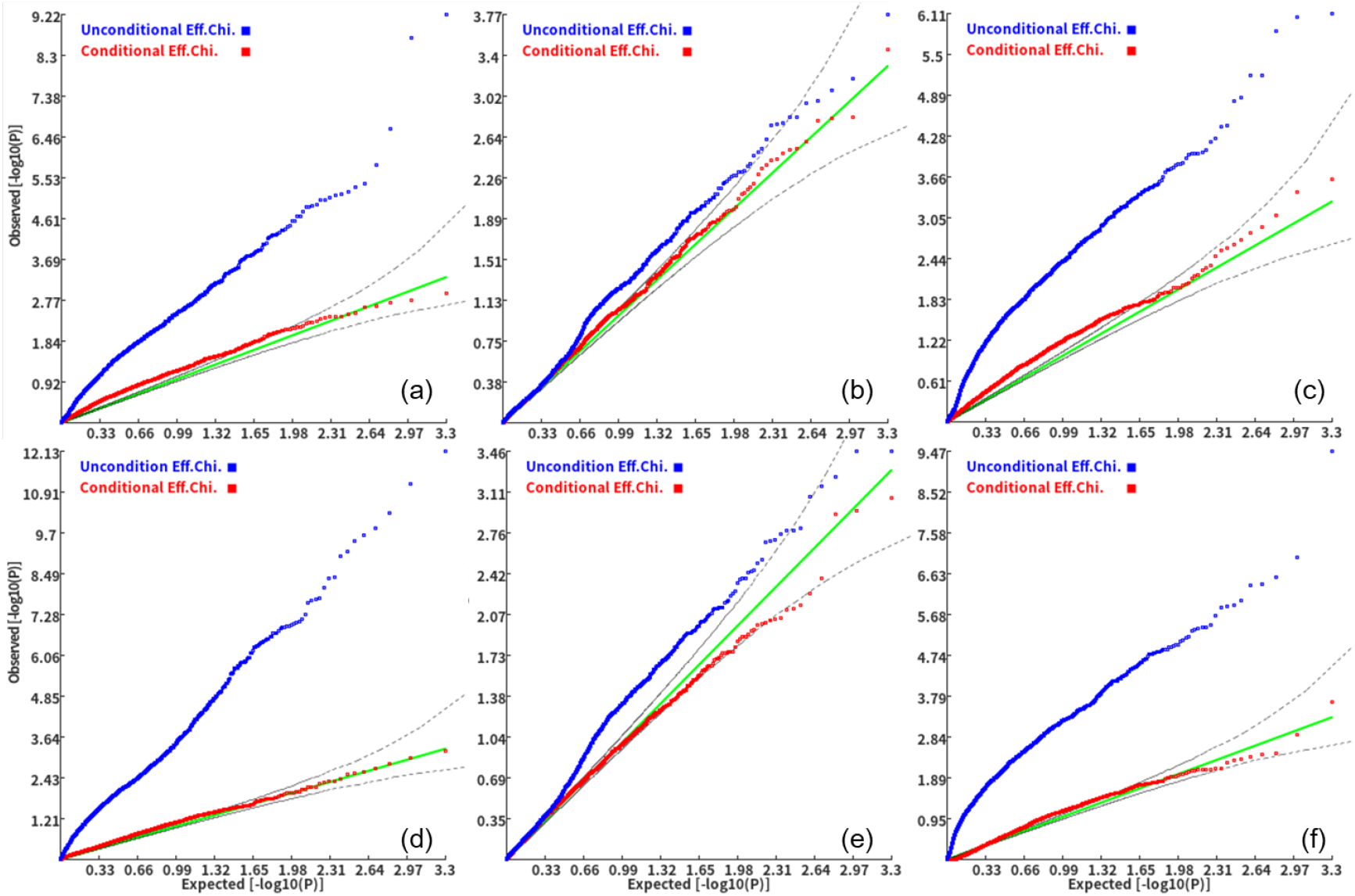
Q-Q plot of conditional gene-based association analysis at different representative gene pairs. **a** and **d**, two genes pairs with similar sizes (SIPA1L2 with 29 variants and LOC729336 with 30 variants), **b** and **e**, two gene pairs with size the first larger than the second (CACHD1 with 41 variants and RAVER2 with 8 variants), **c** and **f**, two gene pairs with size the second larger than the first (LOC647132 with 5 variants and FAM5C with 48 variants). **a, b** and **c**, the formal gene has no QTL and QTL in the latter gene explained 0.5% of heritability. **d, e** and **f**, the formal gene has no QTL and QTL in the latter gene explained 1% of heritability. One thousand phenotype datasets were simulated for each scenario. Unconditional Eff. Chi. (the blue) represents unconditional association analysis at the former gene by the improved test. Conditional Eff. Chi (the red). represents conditional association analysis at the former gene conditioning on the latter gene by the effective chi-squared test.

## Discussion

In the comprehensive and high-resolution landscapes of association between around 1000 phenotypes and more than 20,000 single cell clusters, we found most phenotypes are moderately selectively associated with a fraction of cell types. Specifically, more than 80% phenotypes were selectively associated with less than 10% cell clusters in the above three scRNA-seq datasets. Interestingly, around 20% phenotypes were even selectively associated with less than 1% cell clusters. From the aspect of cell clusters, we observed only a small fraction of cells clusters or cell types were pleiotropic in complex phenotypes. Specifically, only about 10% cell clusters were associated with more than 10% phenotypes, which were named as pleiotropic cell types. Among these pleiotropic cell types, several cell types were related with many phenotypes. For example, fibroblasts in the three single cell datasets were estimated to be involved with more than 40%, 30%, and 20% phenotype sub-categories respectively. This may be because these pleiotropic cell types are prevalent in general biological process. The detailed cell-type specificity of a phenotype and phenotypic pleiotropy of a single cell cluster can be directly retrieved from the website SPA (http://pmglab.top/spa).

In the landscape, we also found significant associations mutually occurred at three blocks (i.e., the CPMP blocks), namely, immune cells – immune phenotypes, brain cells – psychiatry phenotypes and connective tissue, muscle, cardiovascular cells – body measure, musculoskeletal, cardiovascular, metabolism phenotypes (**Fig. 4g**) in the PanglaoDB, which were supported by massive knowledges. Interestingly, very similar enrichment patterns also occurred in the annotation analysis of thousands of cell clusters derived from another independent scRNA-seq database, Human Cell Landscape (**Supplementary Fig. S2a**). Besides testifying the effectiveness of the annotation framework, the repeatable association patterns reaffirmed the hypothesis, at single cell type level, that genes associated with complex phenotypes tend to have selective or preferential expression in phenotype-relevant cell types^8,36^.

Another interesting result was that we could infer reasonably associated cell types for human complex phenotypes even from the usage of mouse scRNA-seq expression profiles. We investigated the performance of annotation in mouse by annotating more than 10,000 cell clusters of mouse from PanglaoDB. Similar to human cells, the three major CPMP blocks were replicated in mouse cells, i.e. immune cells – immune phenotypes, brain cells – psychiatry phenotypes, connective tissue cells, muscle cells, cardiovascular cells – body measure, musculoskeletal, cardiovascular, metabolism phenotypes (**Supplementary Fig. S2b**). This may be regarded as important evidence that genes’ selective expression and their contribution to the development of phenotypes are evolutionarily conserved between human and mouse. This conservation is also an important support for the application of mouse models to study complex phenotypes of human. The robust annotation across species may suggest a practical application, i.e. inferring pathogenic cell types of a complex phenotype by using mouse’s single cell data as proxy. This is because in public domain the available scRNA data of mouse are much more than that of human. Consequently, former may lead to the annotation of phenotype with cell types at higher resolution.

The two application functions will further enhance the practicability of the landscape. To our knowledge, we, for the first time, proposed a method to annotate single cell clusters directly by massive phenotypes, which could gain insights into the possible roles of cell clusters in the development of phenotypes. By annotating a customized cell clusters from scRNA-seq of lungs, we found several cell clusters were significantly associated with respiratory and immune phenotypes sub-categories, which suggested the possible role of them in the development of those phenotypes. In the second application function, thanks to the curated expression profiles of up to 20,000 single cell clusters, we provided accurate identification of phenotype relevant cell types at unprecedented resolution. By identifying the relevant cell types associated with severe COVID-19, we found T cell was a key cell type for the first time from genetic perspective. A recent study reported the weak T cell response in severe patients with COVID-19^34^, which was consistent with our results. Furthermore, the function also prioritized involved genes which could substantially facilitate subsequent investigation of possible pathogenic mechanism of COVID-19.

The sub-categories of phenotypes were a refined classification of phenotypes. Conventionally, diseases and traits were categorized according to their phenotypic characters and the genetic similarity was not considered^37^. Therefore, some phenotypes of the same categories had very different genetic drivers and had few overlapped causal genes, e.g. thyrotoxicosis and adrenal disorder in the endocrine disease category. Thus, traditional phenotypic categories may mislead some studies on the biological mechanism of disease categories. In the present study, we added a new dimension, similarity of associated genes, to further categorize the phenotypes. It should be noted that these associated genes are credibly produced by our selective expression guided conditional association analysis. Based on the sub-categories, we found the association between cell types and phenotypes became more reasonable. We also provided a web-page resource to share these sub-categories. The sub-categories with the supported genes may shed insight into precise diagnosis and even treatment of complex diseases.

The conditional gene-based association test also contributed a lot to the effectiveness of the framework. Due to LD, conventional gene-based association test often detected many indirectly associated genes which would distort the enrichment analysis. Guided by the selective expression of associated genes in prioritized cell types of a phenotype, the conditional gene-based association test effectively purified the list of phenotype-associated genes and led to a more powerful enrichment prioritization of the cell-types in turn^38^. Moreover, we improved conditional gene-based association test by more accurately quantifying correlation of chi-square statistics under alternative hypothesis. It is much more difficult to accurately approximate correlation of chi-square statistics under alternative hypothesis than under null hypothesis^38^. The usage of quantification model for null hypothesis would lead to severe inflation of type 1 error in a conditional gene-based association test^38^, which has been an open question in statistical genetics field. In the present study, we proposed a more accurate model to effectively approximate chi-square statistics correlation under alternative hypothesis which solved the issue of type 1 error inflation in a conditional gene-based association test. The improved gene-based association test with correct type 1 error therefore can detect a purified list of disease-associated genes for the enrichment analysis in annotation.

In conclusion, we provide the most comprehensive, by far, landscape of associations between single-cell clusters and complex diseases and traits by a cross annotation framework. Based on the landscape, phenotypes’ cell-type specificity and cell-types’ pleiotropy in phenotypes are systematically investigated and overviewed. The landscape and the framework may greatly facilitate the cellular-level understandings of tissue specificity and pleiotropy among complex diseases and traits.

## Methods and Materials

### Curation and preprocessing of scRNA-seq data

Three reference scRNA-seq datasets were downloaded from PanglaoDB (human and mouse, https://panglaodb.se/) and HCL (Human cell landscape, http://bis.ziu.edu.cn/HCL/) respectively. The expression data types of scRNA-seq datasets are UMI count or RPKM. The cell clusters information and inferred cell types were also directly downloaded from the corresponding databases. For the scRNA-seq dataset of human lung, we curated the UMI count of single cells derived from normal lung of eight donors in GEO (GSE122960). Then the cells were clustered by Seurat (version 3.1.2).

The expression profiles of cell clusters per dataset were produced according to the following steps. First, the UMI count was converted into the count per million (CPM) to correct for the total number of reads per cell. Second, the expression value was normalized by *log*_2_ (1 + *x*) (*x* is CPM or RPKM, and adding 1 is to avoid the negative infinity) and then the average expression profiles of the cell clusters were calculated. In the mouse scRNA-seq dataset, genes were mapped to their homologous human genes by the R package “biomaRt” (version 2.34.2). Finally, the genes were assigned with HGNC gene symbols and genes without known HGNC gene symbols were removed.

### Curation and preprocessing of GWAS summary statistics of complex diseases and traits

We curated 2250 GWAS summary statistics with full alleles records from Gene Atlas^39^ (http://geneatlas.roslin.ed.ac.uk/), GWAS Atlas^40^ (https://atlas.ctglab.nl/) and Neale Lab UKBB v3 (http://www.nealelab.is/uk-biobank) according to the curation rules of CAUSALdb^41^, which integrated large numbers of publicly available GWAS summary statistics. That is, only GWAS datasets that simultaneously appear in the three databases and CAUSALdb were curated. Therefore, all the 2,250 GWASs could be mapped to CAUSALdb. The population information, sample size, and mapped MeSH term were exacted from CAUSALdb. It should be noted that GWAS Atlas collected the information of GWAS summary statistics from non-UKBB cohort while these datasets actually were provided by other websites, such as GRASP (https://grasp.nhlbi.nih.gov/FullResults.aspx) and PGC^42^ (https://www.med.unc.edu/pgc). Here these datasets were regarded as the GWAS datasets from GWAS Atlas. The curation details of the GWAS datasets were shown in **Supplementary Table S5**. We exacted the p values and chromosome coordinates of all available variants. Non-GRCh37 coordinates were converted to GRCh37 coordinates and the variants that couldn’t be converted were removed. After excluding the GWAS with small sample size (n<10,000), there were 1,871 GWAS summary statistics datasets retained. The GWAS summary statistics of severe COVID-19 was downloaded from http://ftp.ebi.ac.uk/pub/databases/gwas/summary_statistics/GCST90000255/GCST90000255_GRCh38.tsv.gz and the chromosome coordinates were converted to GRCh37 coordinates.

### Selective expression of genes in single-cell clusters

The selective expression of genes was calculated by the robust-regression z-score method (REZ, http://grass.cgs.hku.hk/limx/rez/). For the three reference scRNA-seq datasets, we directly calculated the selective expression with the average expression of the clusters. For the scRNA-seq dataset of human lung, we combined average expression of cell clusters with the average expression of the reference cell clusters (PanglaoDB’ s human dataset). We then calculated the selective expression of lung cell clusters in the combined expression profiles.

### The improved conditional gene-based association analysis

In the iterative prioritization procedure of SPA, we improved and extended our previous effective chi-square test^38^ to perform the conditional gene-based association analysis. The extension was proposed to be a more effective model and then remove redundant association signals. The first step of the analysis is to produce effective chi-square statistics for genes. Suppose there are in total *n* loci in a set of genes and *m* loci in its subset of genes to be conditioned. Each locus has a p-value for phenotype association from a GWAS. The p-value is converted to corresponding chi-square statistics with degree of freedom 1. According to Li et al.^38^, each locus is assumed to have a virtually independent chi-square statistics which additively contributes to the observed marginal chi-square statistics. Here the correlation of chi-square statistics between two loci is found to be approximated by the absolute value of genotypic correlation |**r**| when the expected chi-square statistics is large under alternative hypothesis (See derivation in **Supplementary Note 1**), which was an unsolved issue in Li et al.^38^. According to Li et al.^38^, under null hypothesis the correlation of two chi-square statistics is equal to *r^2^*. Therefore, the n loci are separated into two groups. Group 1 contains variants with p-values ≤ 0.05 and group 2 contains the rest. For n1 variants in group 1, virtual independent chi-square statistics 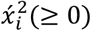 and degree of freedom *d_i_*(>0) are calculated by the following formula,

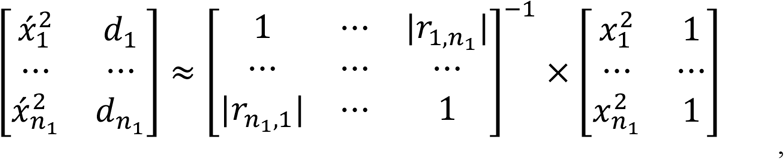

where 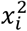 is the observed chi-square statistics and |*r_i,j_*| is the absolute value of LD correlation coefficient.

The effective chi-square statistics *Ś*_*g*1_ with degree of freedom *d*_*g*1_ for group 1 is then obtained by:

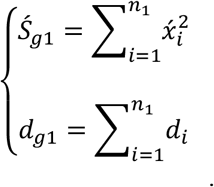

For group 2, the effective chi-square statistics is then also calculated by the same formula except that the |*r_i,j_*| is replaced with 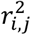. As the virtually independent chi-square statistics are expected to be larger than 0, the R package NNLM (https://rdrr.io/cran/NNLM/) was used to calculate the non-negative effective chi-square statistics and degree of freedom. The effective chi-square statistics *ś_n_* and degree of freedom *d_n_* of the n loci are obtained by adding up the effective chi-square statistics and degree of freedom of the two groups. The effective chi-square statistics *ś_m_* and degree of freedom *d_m_* of the m loci to be conditioned are calculated by the same formula and procedure. The conditional gene-based association test is then obtained according to conditionally effective chi-square statistics formula,

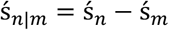

and degree of freedom

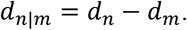

In a GWAS, all genes are first calculated with the unconditional gene-based association p-values by the above effective chi-square statistics under null hypothesis. Given a p-value cutoff, the significant genes are extracted and subject to the conditional gene-based association analysis. When there are multiple significant genes in a LD block, the genes are conditioned one by one in a pre-defined order. Here we suggest assigning the genes within 5 MB into the same LD block. The conditional p-value of the first gene in the order is defined as its unconditional p-value. The conditional p-value of the second gene is obtained by conditioning on the first gene; and that of the third gene is obtained by conditioning on the top two genes. The conditional p-values of subsequent genes are obtained according to the same procedure. In the first iteration, the genes’ order is defined according to genes’ unconditional p-values. Since the second iteration, the order is defined according to genes’ selective expression in the prioritized cell types^8^ by Wilcoxon rank-sum test. The whole analysis procedure is implemented in KGGSEE (http://pmglab.top/kggsee).

### Association of a single-cell cluster with a phenotype by selective expression enrichment analysis

In the iterative prioritization procedure of SPA, the other test is whether a single-cell cluster is associated with a phenotype according to selective expression enrichment of phenotype-associated genes. The significantly associated genes of a phenotype are assigned into one group; all other genes on the genome are put into the other group. The Wilcoxon rank-sum test was then adopted to compare whether the former is enriched with higher selective expression than the latter in a tested cell cluster. The rank-sum test is carried out at all the cell cluster-phenotype pairs.

### Gene-based association analysis with GWAS p-values

The variants in each downloaded GWAS dataset were mapped onto genes according to their physical positions on chromosomes. To cover more regulatory variants, variants within 5kb extended regions of gene boundaries at both sides were included. The gene boundaries in RefGene and GECODE were adopted. Ancestry-matched genotypes from 1000 Genomes Project were used to account for dependency of association signals in the conditional and unconditional gene-based association tests. All the analyses were carried out through our software tool KGGSEE.

### Classification of phenotypes

Firstly, according to the corresponding MeSH terms mapped by CAUSALdb^41^, phenotypes were manually classified into 19 clinical categories. Then the phenotypes were clustered by the associated genes within each of the clinical category by the following method.

Suppose for a phenotype *p*, its associated genes were labeled as *G_p_* = (*g*_*p*1_, *g*_*p*2_,…, *g_pn_*). The union of all phenotypes’ associated genes was labeled as *G* = (*g*_1_, *g*_2_,…, *g_n_*). We mapped the *G_p_* to *G* and got the n-dimensional binary vector *GV_p_* = (1, 0,…, 1) (1 represents mapped gene and 0 represents unmapped gene). Then we clustered the phenotypes of each clinical category by the hierarchical clustering with the n-dimensional binary vector. The Euclidean metric was used in the hierarchical clustering and the clustering threshold was empirically set as 20.

Finally, the phenotypes were classified into 107 sub-categories by combining the clinical categories with the above clustering results. The naming convention of sub-category was “clinical category name” + “_” + “cluster id”. For example, the sub-category “musculoskeletal_1” means the cluster “1” in the clinical category “musculoskeletal”.

### Similarity measure of phenotype categories

For phenotype categories, say *a* and *b*, their phenotype sets were *P_a_* = (*x*_1_, *x*_2_,…, *x_m_*) and *P_b_* = (*y*_1_, *y*_2_,…, *y_n_*) respectively. Then the similarity of *a* and *b* was measured by the averaged Jaccard coefficients of each phenotype in *P_a_* and each phenotype in *P_b_* according to the associated genes. If phenotype *a* and *b* were same, the Jaccard coefficients of same phenotype pairs were excluded for averaging. The formula for the similarity measure is shown below.

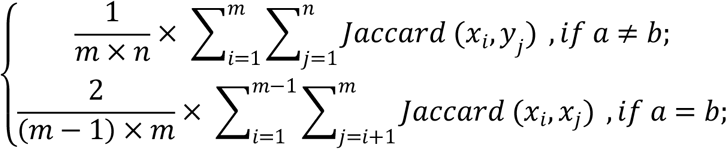

### Enrichment analysis of phenotype sub-categories and inferred cell types, source tissues or cell clusters

Assumed that the association profile contains *N* phenotypes and *M* cell clusters, *k* pairs of cluster-phenotype were significantly associated by the FDR correction. For the phenotype sub-category *a*, its phenotypes were *P_a_* = (*p*_1_, *p*_2_,…, *p_n_*), *P* represents a phenotype. For the inferred cell type, source tissues or cell cluster *u*, single cell clusters belonging to *u* are labeled as *C_u_* = (*c*_1_, *c*_2_,…, *c_m_*), *c* represents a single cell cluster. Particularly, when *u* represents a cell cluster, *C_u_* only contains itself, *j* pairs between phenotypes in *P_a_* and clusters in *C_u_* are significantly associated. Then the enrichment p values of *a* and *u* are calculated by the hypergeometric test with following formula.

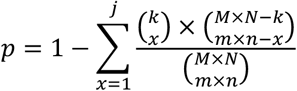

### Simulations for investigating type 1 error and power of the conditional gene-based association test

Extensive computer simulations were performed to investigate type 1 error and power of the conditional gene-based association test. To approach the association redundancy pattern in realistic scenarios, we used real genotypes and simulated phenotypes. The high-quality genotypes of 2,507 Chinese subjects from a GWAS^43^ were used and phenotypes of subjects were simulated according to the genotypes under an additive model. Given total variance explained by *n* independent variants, V_g_, the effect of an allele at a bi-allelic variant was calculated by 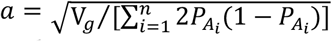, where *P_A_i__* was the frequency of alternative alleles. The total expected effect *A* of a subject was equal to a*[the number of alternative alleles of all the n variants]. The phenotype of each subject was simulated by *P*=A+e, where *e* was sampled from a normal distribution *N*(0, 1-V_g_). We randomly sampled three pairs of genes, i.e. SIPA1L2 vs. LOC729336, CACHD1 vs. RAVER2, and LOC647132 vs. FAM5C. They represented three scenarios where the nearby gene (i.e., the first gene) had more similar (SIPA1L2 vs. LOC729336), larger (CACHD1 vs. RAVER2) and smaller (LOC647132 vs. FAM5C) size than the target gene (i.e., the second gene) respectively in terms of SNP number. In the investigation of type 1 error, the target gene had no QTLs while the nearby gene had one or two QTLs. In the investigation of the statistical power, both genes had QTLs.

For the purpose of power comparison, the likelihood ratio test based on linear regression was adopted to perform the conditional gene-based association analysis with raw genotypes. In the full model, genotypes of all SNPs encoded as 0, 1, or 2 according to the number of alternative variants entered the regression model as explanatory variables. In subset model, the SNPs of the nearby genes entered the regression model. The calculation of the likelihood ratio test was performed according to conventional procedure. The R packaged “lmtest” (version 0.9.37) was adopted to perform the likelihood ratio test.

## Description of Supplemental Data

Figures S1–S4, Tables S1–S8, Note 1.

## Acknowledgements

We thank The PanglaoDB, Human Cell Landscape, to provide access for scRNA-seq expression data. We also appreciate GWAS Atlas, Gene Atlas, Neale lab, CAUSALdb for providing the GWAS summary statistics data and phenotype meta information.

## Availability of data and materials

The SPA web application is available at http://pmglab.top/spa. The KGGSEE (V1.0) is available from http://pmglab.top/kggsee. Furthermore, the source code of KGGSEE (V1.0) can be downloaded from Github (https://github.com/pmglab/KGGSEE). The code is released under MIT license (https://opensource.org/licenses/MIT). The scRNA-seq expression dataset of PanglaoDB was obtained from https://panglaodb.se (prior to February 2020) and Human Cell Landscape from http://bis.ziu.edu.cn/HCL (prior to April 2020). The human lung scRNA-seq expression dataset was curated from GEO (GSE122960). The GWAS summary statistics dataset of Gene Atlas was obtained from http://geneatlas.roslin.ed.ac.uk, GWAS Atlas from https://atlas.ctglab.nl, Neale Lab UKBB v3 from http://www.nealelab.is/uk-biobank and COVID-19 from http://ftp.ebi.ac.uk/pub/databases/gwas/summary_statistics/GCST90000255/GCST90000255_GRCh38.tsv.gz.

## Ethics approval and consent to participate

Not applicable.

## Consent for publication

Not applicable.

